# Distinct profiles of spatio-temporal brain dynamics along symptoms dimensions in autism

**DOI:** 10.1101/2021.03.11.434353

**Authors:** Emeline Mullier, Nada Kojovic, Solange Denervaud, Jakub Vohryzek, Patric Hagmann, Marie Schaer

## Abstract

Autism Spectrum Disorders are accompanied by atypical brain activity and impairments in brain connectivity. In particular, dynamic functional connectivity approaches highlighted aberrant brain fluctuations at rest in individuals with autism compared to a group composed of typically developed individuals, matched in age and gender. However, the characterization of these variations remains unclear. Here, we quantified the spatio-temporal network dynamics using two novel dynamic group-based measures, namely system diversity and spatio-temporal diversity. Using the public database ABIDE 1, we explored the differences between individuals with autism and typically developed individuals. Our results show evidence that individuals with autism have atypical connectivity patterns over time characterized by a lower integration of heterogeneous cognitive processes and unstable functional activity, except for the default mode network presenting its own specific dynamic pattern. Within the autism group, we find this pattern of results to be stronger in more severely affected patients with a predominance of symptoms in the social affect domain. However, patients with prominently restricted and repetitive behaviours demonstrate a more conservative profile of brain dynamics characterized by a lower spatio-temporal diversity of the default mode network.

## 1. INTRODUCTION

Autism Spectrum Disorder (ASD) is a group of neurodevelopmental disorders characterized by deficits in social communication and the presence of repetitive and restricted behaviours (RRB) (Association American Psychiatric 2013). Difficulties with transitions and change, insistence on sameness, and preference for stable, repetitive patterns of activities are among the disorder’s core features. These behavioral patterns are often linked with specific executive functioning profile characterized by reduced cognitive flexibility (Corbett et al. 2009; Dajani and Uddin 2015) associated with the severity of RRB symptoms (Lopez et al. 2005). In social domain, cognitive inflexibility might also impair the way affected individuals switch rapidly their focus of interest in highly dynamic social interactions, making these interactions challenging to understand with a possibility of causing social anxiety. Thus, these two symptoms’ dimensions, interactions deficits on one hand and the repetitive and restricted behaviors on the other, quite different in their manifestations, might indeed have a common causal pathway.

A growing body of research showed that individuals with ASD exhibit atypical brain connectivity profiles characterized by both insufficient and excessive connectivity [For review (Hull et al. 2017)]. These disruptions in neural network organization (Müller and Fishman 2018; Morgan et al. 2018) reflect atypical underlying neural processes (Jones et al. 2018). Connectivity studies in ASD population highlighted impaired connectivity within the salience network (SAL) This feature allowed an accurate classification between typically developed children and children with ASD and was a reliable predictor of RRB scores (Uddin et al. 2013). Aberrant connectivity patterns have also been consistently reported in other long-range functional networks namely the default mode network (DM), the frontoparietal (FP) (Abbott et al. 2016), the sensorimotor (SM) and the visual (VIS) networks (Uddin 2015).

Beyond the connectivity profiles of the functional systems, their dynamic interplay over time is essential to understand how they support complex cognitive processes. The coupling of the FP and the DM networks, orchestrated by the SAL network, is central to the neural dynamics underlying cognitive flexibility. The SAL network acts as a fast integrator and regulator of the FP and the DM networks (Menon and Uddin 2010) to adapt behaviour through a twofold mechanism: (i) fast and automatic perceptive bottom-up signalling and (ii) a high-order system related to context specific cues enhancing access to goal directed actions (Menon 2015). Consequently, salience network control impacts attentional and executive abilities (Taghia et al. 2018). Generally, alterations observed in these three core networks could lead to more cognitive inflexibility (Uddin, Supekar, and Menon 2010).

Recent dynamic functional connectivity approaches (Preti, Bolton, and Van De Ville 2017) showed that individuals with ASD present local aberrant brain fluctuations at rest (Uddin 2021). A few studies showed altered dynamics in several core networks, such as the FP network (Kupis et al. 2020; Guo et al. 2019; 2020) [i.e. the anterior cingulate cortex (Guo et al. 2020)], hubs of the DM network [prefrontal and posterior cingulate cortices], the thalamus or some sensorimotor regions (Fu et al. 2019; Guo et al. 2020). The connectivity states were also characterized by less frequent transitions between salience-central executive and DM-medial frontoparietal in children with ASD (Marshall et al. 2020) and by fewer transitions to an unstable states in adults with ASD, negatively correlated with the gold standard Autism Diagnosis Observation Schedule ADOS score (Watanabe and Rees 2017). Another study from Harlalka and al. (Harlalka et al. 2019) also showed a positive correlation between symptoms severity and reduced flexibility.

A joined effort represented in form of large and publicly available neuroimaging datasets is necessary to address the tremendous heterogeneity and complexity of ASD. The Autism Brain Imaging Data Exchange (ABIDE) project (Di Martino et al. 2014; Di Martino et al. 2017) promotes data sharing on ASD MRI cohorts across sites. Using this rich resource, it has been shown that the degree of variability in connection strength across time within the cingulate, temporal and parietal lobes was positively correlated with symptoms severity (Harlalka et al. 2019; Li et al. 2020).

Furthermore, the decreased variability in ASD negatively correlates with social motivation and social relating (He et al. 2018). Moreover, longer dwell times in weak states of functional connectivity found in ASD (Yao et al. 2016) were associated to higher levels of autistic traits (Rashid et al. 2018). In the study of Watanabe and al. (Watanabe and Rees 2017), IQ scores of the group of typically developed individuals were predicted by the neural transition frequency while IQ scores of ASD group were rather predicted by the stability of their brain dynamics. These findings are the first attempts to bridge clinical observations to neural network dynamics.

Based on the principal of functional co-activation patterns (Tagliazucchi et al. 2012; Liu et al. 2018), spatio-temporal connectomics approach aims to capture the propagation of functional connectivity on the structural connectome by encoding transient network of spatio-temporal activity (Griffa et al. 2017). Two novel dynamics measures, namely system diversity (SD) and spatio-temporal diversity (STD), have been developed to help characterizing this spatio-temporal activity in terms of its diversity of recruitment of cognitive systems (SD) and its spatial self-similarity over time (STD) (Vohryzek et al. 2019). SD and STD were shown to capture an increased integration of various functional systems with age along with a higher self-similarity between activity patterns over time suggesting a maturation of the repertoire of integrated spatio-temporal pattern with age (Vohryzek et al. 2019), and positively influenced by experience such as schooling (Denervaud et al., Under review). Despite the evidence of altered brain fluctuations at rest in individuals with ASD, fundamental brain mechanisms reflected by the reorganization of the spatio-temporal activity patterns remain poorly understood, as well as their impact on clinical outcomes. The measures of SD and SRD thereby represent a promising first step toward the investigation of combined structural and functional coupling alterations.

Taken together, the current findings converge on the idea of disrupted brain dynamics in individuals with ASD compared to typically developed individuals (TD). However, the disruptions of spatio-temporal dynamics in individuals remain unclear. Besides, it is unknown how the underlying altered dynamic translates to the heterogeneity of profiles both in terms of the symptoms severity but also the preponderance of symptoms along the two main symptom dimensions: social communication and interaction deficits (social affect - SA) and RRB. The purpose of this study was to explore these spatio-temporal dynamics according to these distinct profiles of symptoms with different levels of severity in SA and/or RRB.

Here, we applied these two novel measures, SD and STD, on 185 individuals with ASD (age 12.5±2.9) matched with 185 typically developed individuals (age 12.8±2.6) issued from ABIDE 1 to investigate the temporal reorganization of brain activity over time. We investigated these changes in the whole brain and in different functional systems to identify which systems would be affected in the ASD group. Besides, we examined how these measures vary along the symptoms dimensions by splitting the individuals with ASD into different symptoms profiles based on two core symptomsmeasures with ADOS: SA and RRB with: i) severe vs moderate overall level of autistic symptoms; ii) severe vs moderate level of social deficits and iii) severe vs moderate level of RRB. We expect that different spatio-temporal patterns may be related to the different symptoms domains in ASD. We hypothesize that the predominant social affect deficits could be associated to hypervariability of brain activity at rest while the severity of RRB could be the result of disrupted cognitive flexibility mechanisms.

## 2. MATERIAL AND METHODS

### 2.1 Participants

ABIDE 1 preprocessed database (Craddock 2013) includes MRI images from 539 individuals with ASD and 537 typically developed individuals with their corresponding phenotypic information from 20 different sites. A direct measure of autistic symptoms was obtained for the participants using the Autism Diagnostic Observation Schedule 2^nd^ Edition (ADOS-2), Module 3 (Lord and Jones 2012). To target the pure extent of symptoms of autism relatively independent of the participants’ characteristics such as age or verbal functioning, we used the Comparison Severity Scores-CCS ranging from 1 to 10 (Gotham, Pickles, and Lord 2009). Additional to the comparison severity scores that indicate the overall severity of autistic symptoms we also included the severity scores across domains of Social Affect (SA) and Restricted and Repetitive Behaviours (RRB). This permitted a more precise insight on the separate dimension of autistic symptoms (Hus, Gotham, and Lord 2014).

For this study, inclusion criteria for subjects were i) low motion during BOLD acquisition (framewise displacement values FD<.5 mm); ii) age of the subject < 30 years old; iii) direct measure of autistic symptoms obtained using ADOS module 3 and iv) availability of Comparison Severity Scores – CSS: total, social affect (SA) and repetitive and restricted behaviors (RRB). Furthermore, the inclusion criteria for sites were a minimal number of 30 subjects (TD and ASD taken together). According to these criteria, 185 patients from 14 different sites were included in the analyses. A matched TD group was created based on average demographic values obtained from the ASD group (Figure 1A).

**Figure 1:**
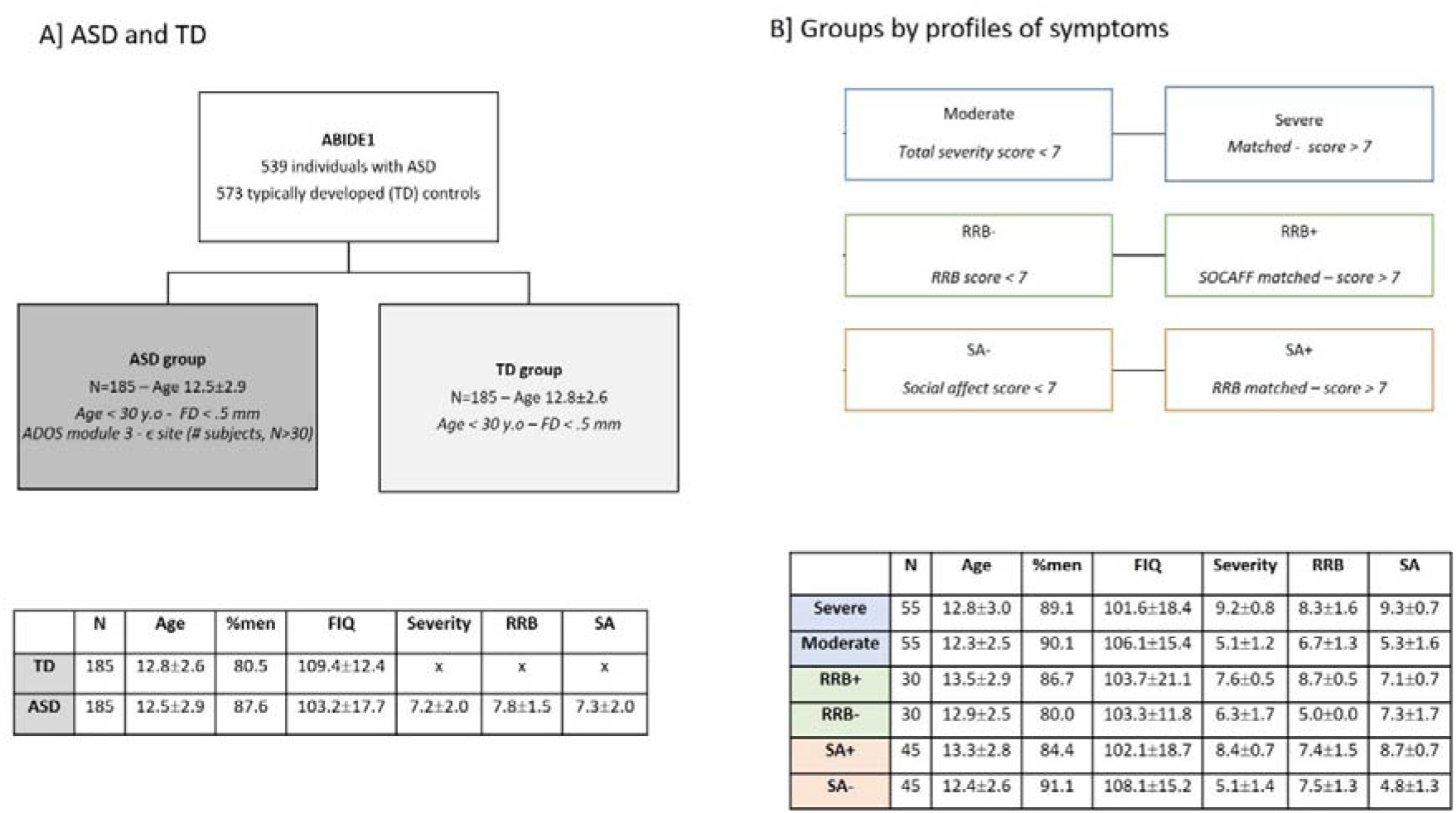
Flowcharts presenting the criteria and demographic information for the selected subjects for our different analyses. (A) From the original 539 individuals with ASD and 573 controls, data were discarded based on the subject’s age and the quality of the data. (B) The ASD group is subdivided into different subgroups based on different profiles of symptoms based on three ADOS derived scores (blue) their total severity score, (orange) their social affect score and (green) their RRB scores.

### 2.2 MRI acquisition and processing

Acquisition details for each site can be found on ABIDE website (http://fcon_1000.projects.nitrc.org/indi/abide/). The pre-processed resting-state functional MRI provided in ABIDE 1 in MNI space were directly downloaded from ABIDE 1 website (http://preprocessed-connectomes-project.org/abide/) using the Connectome Computation System pre-processing strategy with bandpass filtering and without global signal regression. Details about the pipeline can be found here (http://preprocessed-connectomes-project.org/abide/ccs.html). The preprocessed fMRI signals were averaged for each region of multiscale Lausanne 2018 atlas scale 4, comprising 506 brain regions (448 cortical regions) in MNI space and available in the ConnectomeMapper3 (https://doi.org/10.5281/zenodo.3514694).

### 2.3 System Diversity (SD) and Spatio-Temporal Diversity (STD)

The fMRI time series were processed using the spatio-temporal connectome (Griffa et al. 2017). The corresponding codes are available in (https://github.com/agriffa/STConn). A template average structural connectome was used as the structural connectivity constraint for the construction of the multilayer spatio-temporal connectome (https://doi.org/10.5281/zenodo.2872624). For the two groups ASD and CTRL, a series of weakly connected components (CCs) was thus generated. From these series of CCs, SD and STD were estimated (Vohryzek et al. 2019) (https://github.com/ivohryzek/STconnectomics_dvlp) at the global scale and at the functional system scale. A summary of the pipeline is presented in (Figure 2).

**Figure 2:**
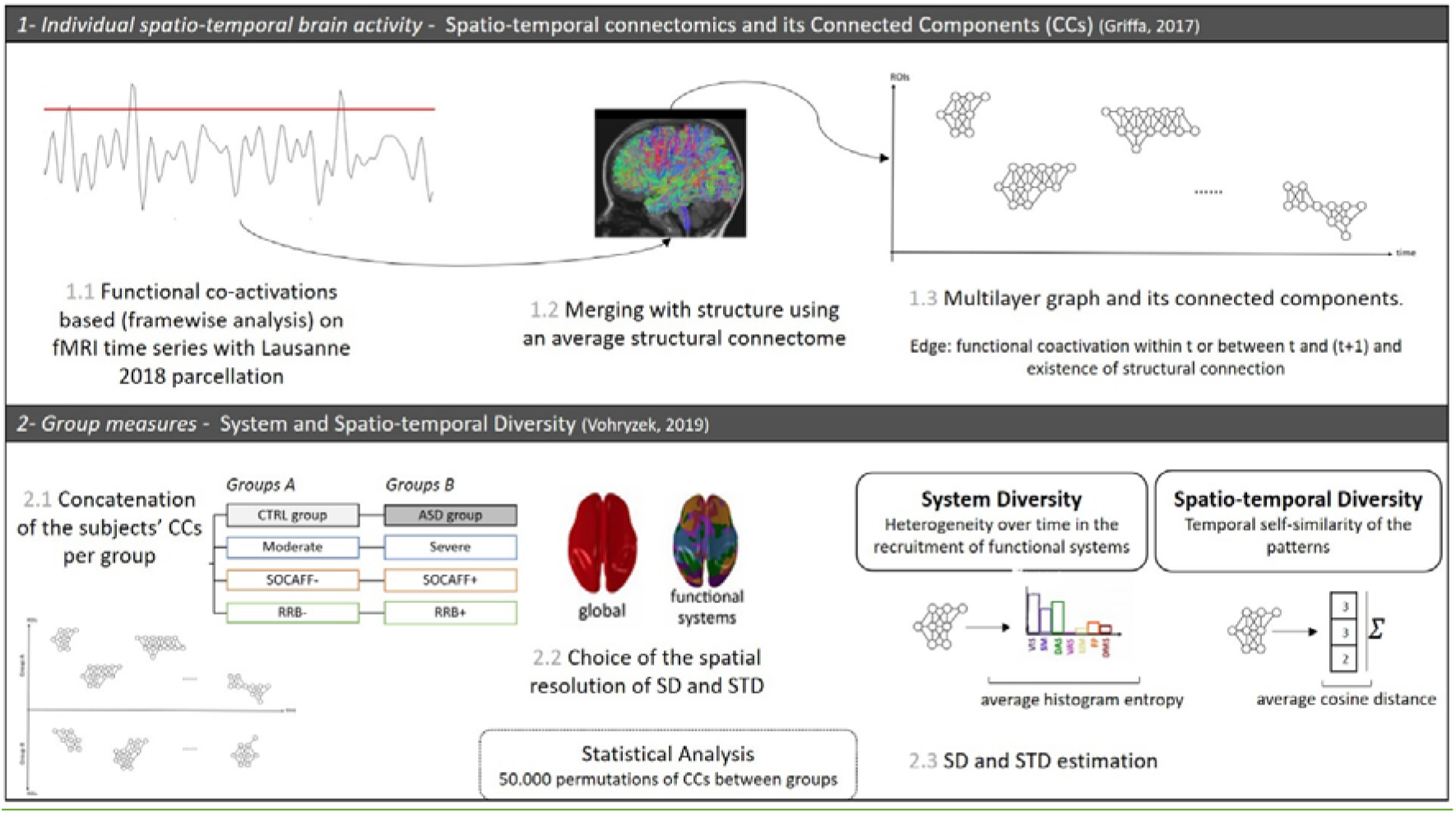
Summary of the pipeline (Adapted from (<u>Griffa et al. 2017</u>) and (Vohryzek et al. 2019)). [1] Main steps of the spatio-temporal connectomics approach. Point process analysis (Tagliazucchi et al. 2012) was applied to the fMRI time series based on Lausanne 2018 parcellation. From these activations, we build a multilayer graph in which an edge was created (i) if there was a functional co-activation within the same time point and a structural connection in the average structural connectome (ii) and if there was a functional co-activation between two successive time points and a structural connection. [2] From the individual multilayer graphs, we extracted the weak connected components of the subjects. The connected components were concatenated by group for group comparisons [2.1]. We computed SD and STD at two different spatial scales: (i) the global scale, meaning that we considered the entirety of the CCs (ii) the functional systems’ scale: SD and STD were estimated for each system and each group considering the CCs belonging to the corresponding group and with at least 20% of its regions attributed to this specific functional system.

### 2.4 Group analyses

We compared changes in SD and STD for different groups. First, we investigated the differences in these measures between the TD group and the ASD group. Then, we performed three analyses along the symptoms’ dimensions. To discern the symptom heterogeneity, we divided the ASD sample into three different symptoms profiles. The division was performed as follows: (i) ASD with moderate symptoms [Moderate, CCS<7] vs ASD with more severe symptoms [Severe, CCS>7] based on their total severity score (ii) ASD with more severe RRB score [RRB+, CCS_RRB>7_] vs ASD with less severe RRB [RRB-, CCS_RRB<7_] for matched SA scores and, (iii) ASD with more severe social affect score [SA+] vs ASD with less severe social affect score [SA-] for matched RRB scores (Figure 1, right panel). For each group, we computed a value of SD and STD at the global scale (whole-brain level) and at the functional systems level, meaning of SD and STD value for each of the seven functional systems from Yeo’s parcellation(Yeo et al. 2011). P-values were assessed based on the distribution of randomized SD and STD values with 50 000 permutations of subjects between the two groups. At the functional systems scale, results were corrected for multiple comparisons using Bonferroni correction. More specifically, we considered a p-value as significant if *p* <.007 (which corresponds to the usual Significant level of .05 divided by the number of functional systems i.e., seven).

## 3. RESULTS

### 3.1 Spatio-temporal diversity and System diversity in individuals with ASD compared to TD group

We estimated SD and STD at two different spatial scales: 1) the global scale (whole-brain level); and 2) the functional systems scale. At the global scale, no significant changes were observed between ASD and TD (Figure 3, top panels). At the functional systems scale, the differences in SD and STD showed heterogeneous patterns. First, SD was lower in the visual network (*p*=.003), while higher in the DM (*p*=.004) for the ASD group compared to the TD group. Second, STD was lower in the DM (*p*<.001), while higher in the limbic system (*p*=.001), the dorsal attention network (*p*=.002) and the visual system (*p*=.006) for the ASD group compared to the TD group (Figure 3, bottom panels).

**Figure 3:**
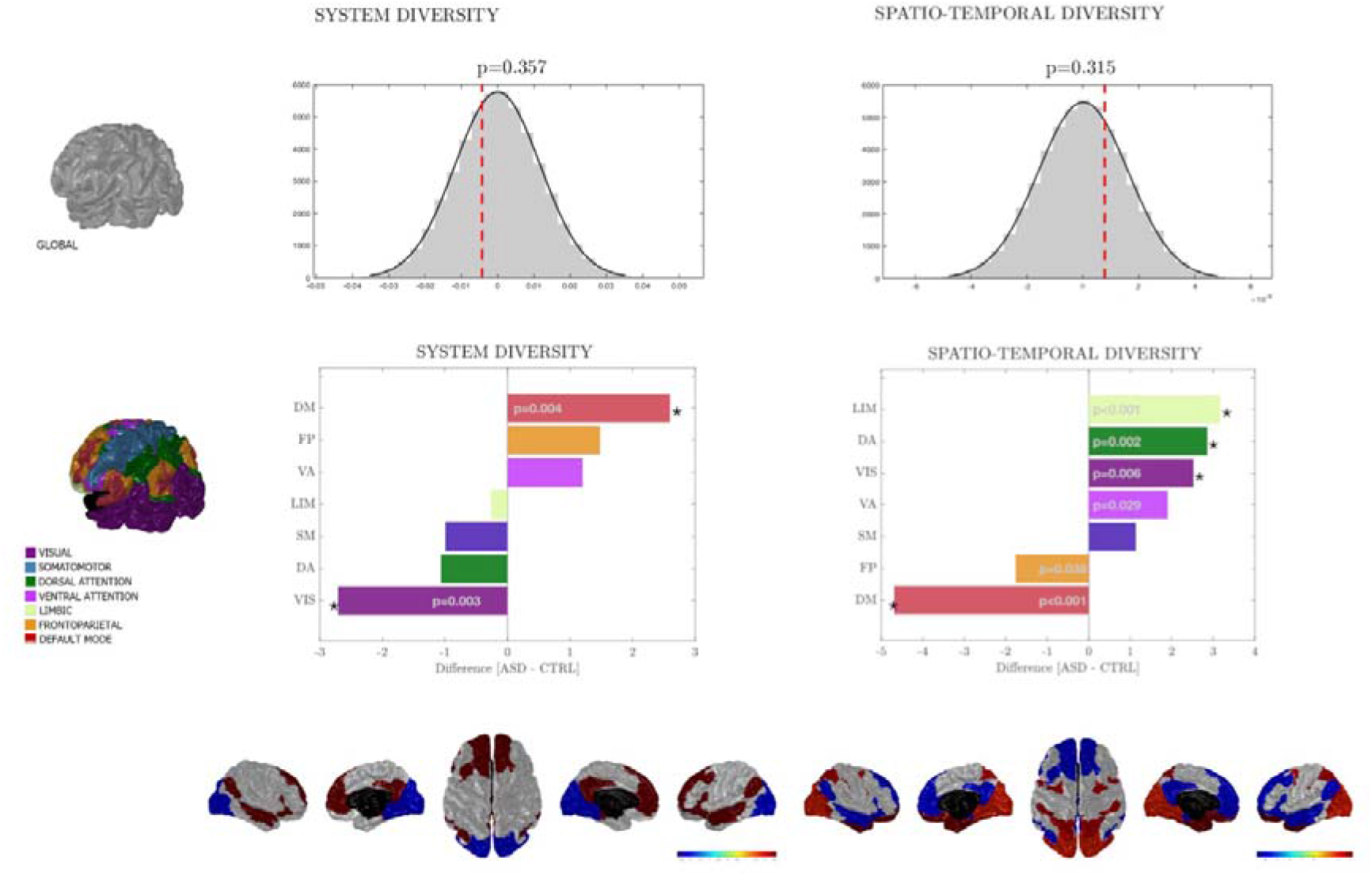
(Top) Difference (ASD-TD) at the global scoleforSD (left) and STD (right). The grey distribution are rondom volues after 50 000 permutations of subjects between groups. The red dot line corresponds to the difference between the two groups ASD and TD. (Middle, bottom) Difference (ASD-TD) at the functional systems scale for SD (left) and STD (right). The black asterisks indicate significance after correction for multiple comparison with Bonferroni (p<.007). P-values were estimated based on a randomized distribution after 50 000 permutations of subjects between groups. (Bottom) Representation of the brain surface of the significant z-scored differences between SD and STD for the functional systems.

### 3.2 Influence of severity of symptoms

To evaluate the extent to which these altered dynamics relate to the severity of symptoms, we repeated the previous analysis with the different subgroups of different symptoms profiles: (i) severely affected [severe] and moderately affected [moderate]; (ii) predominant social affect symptoms [SA+] and moderate social affect symptoms [SA-]; as well as (iii) predominant RRB symptoms [RRB+] and moderate RRB symptoms [RRB-].

#### 3.2.1 Total severity: severe and moderate

At the global scale, no significant differences between the ASD group with more severe levels of symptoms [severe] and the group with moderate levels of symptoms [moderate] were observed in the ASD group (Figure 4, left). However, at the functional systems level, results were similar across the different systems. First, SD was lower in the sensorimotor network (*p*=.006) for the severe symptoms group compared to the moderate symptoms one (Figure 4, right). Second, STD was higher in the DM, the sensorimotor network, the dorsal and ventral attention networks (p<.001 for all of them) and the frontoparietal network (*p*=.001) for the severe group compared to the moderate one.

**Figure 4:**
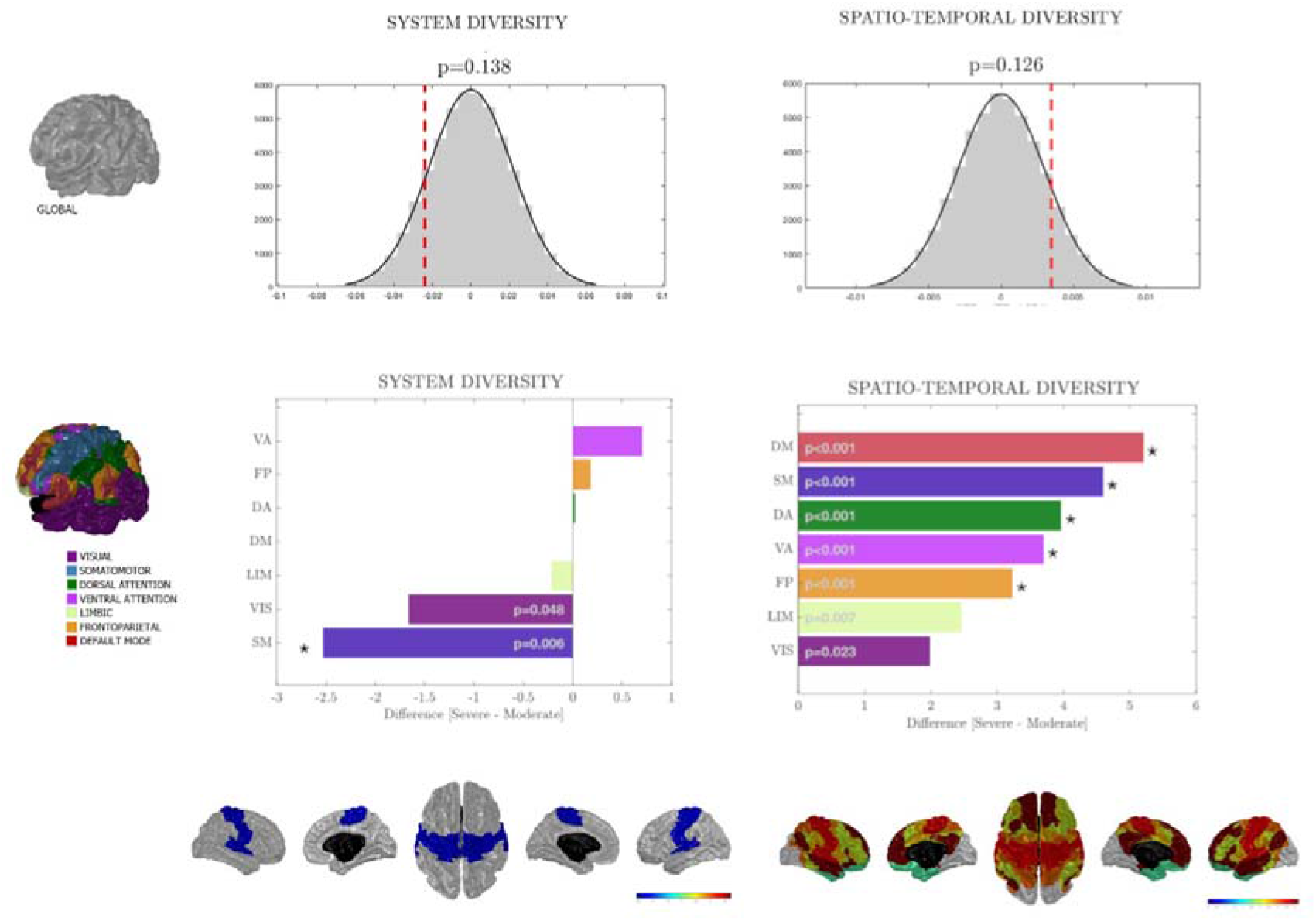
(Top) Difference between the group more severely affected [severe] and the group moderately affected [moderate] (severe - moderate) at the global scale for SD (left) and STD (right). The grey distribution are random values after 50 000 permutations of subjects between groups. The red dot line corresponds to the difference between the two groups severe and moderate. (Middle, bottom) Difference (severe - moderate) at the functional systems scale for SD (left) and STD (right). The black asterisks indicate significance after correction for multiple comparison with Bonferroni (p<.007). P-values were estimated based on a randomized distribution after 50 000 permutations of subjects between groups. (Bottom) Representation of the brain surface of the significant z-scored differences between SD and STD for the functional systems.

#### 3.2.2 Social Affect and Restricted and Repetitive Behaviours

At the global scale, SD was significantly lower in the SA+ group than in the SA-group. However, no significant differences between the RRB+ and RRB-groups were observed in SD (*p*=.101), nor STD (*p*=.430). At the functional systems level, both SD and STD were affected by the levels of symptoms across the social affect and RRB domains. First, SD was lower in the visual and sensorimotor systems for patients with more severe symptoms in social affect domain SA+ (*p*<.001 and p=.001 respectively) (Figure 5, left top panel). Second, for the SA+ group, STD was higher in the sensorimotor cortex and the ventral attention system (*p*<.001 for both) but also within the DM network (*p*=.002). Contrarily, STD for the RRB+ group compared to the RRB-group was lower in the DM network (*p*<.001).

**Figure 5:**
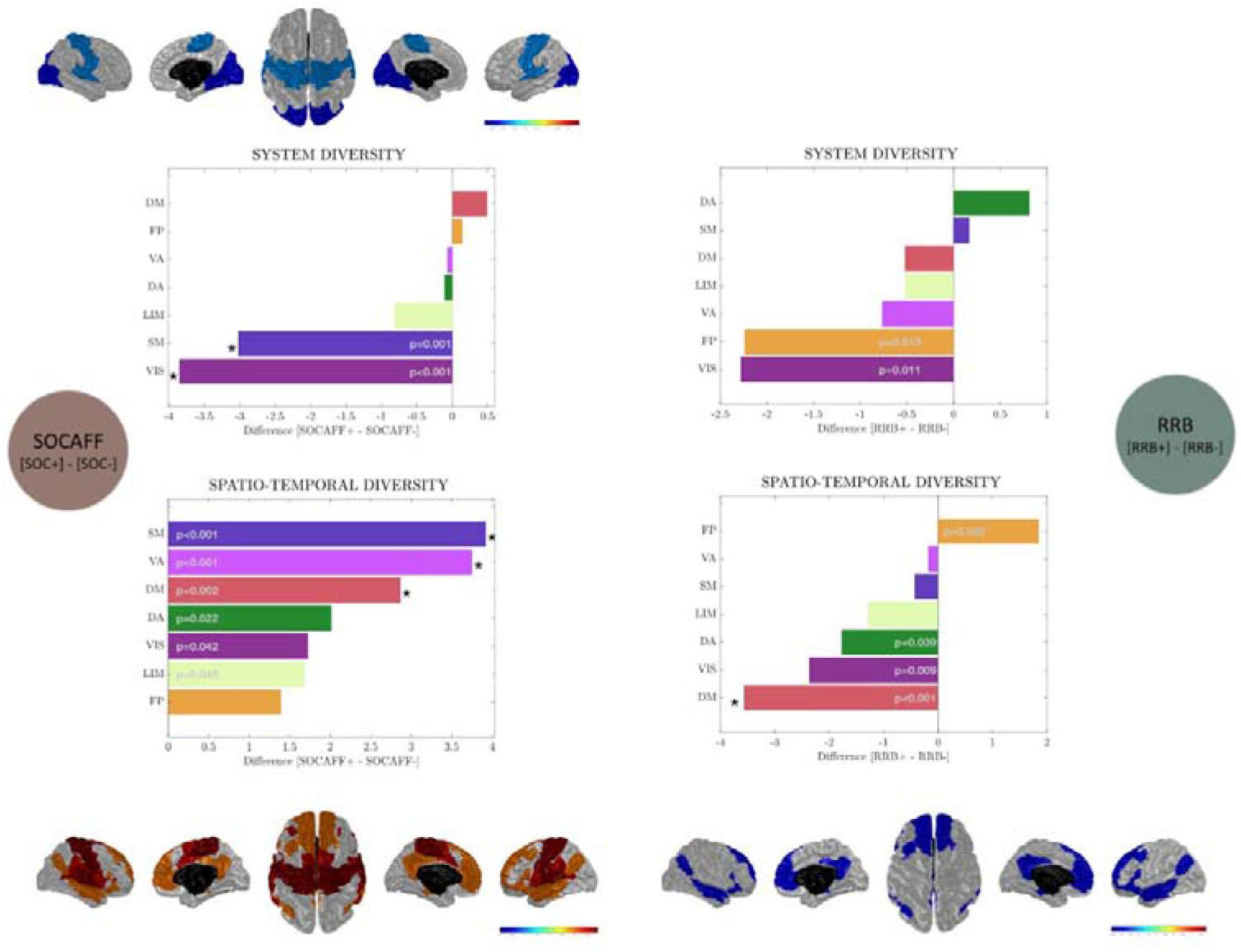
(Left) z-score difference (SA+ - SA-)for SD (top) and STD. The colormap correspond to the value of the differences. Only the systems with significant differences after correction for multiple comparison (p<.007, 50 000 permutations) are coloured. (Right) z-score difference (RRB+ - RRB-)for SD (top) and STD. The colormap correspond to the value of the differences. Only the systems with significant differences after correction for multiple comparison (p<.007, 50 000 permutations) are coloured.

## 4. DISCUSSION

Given the importance of spatio-temporal brain dynamics for cognitive processes, we studied the differences in SD and STD measures between a group of children and adolescents with ASD and their age-matched TD peers at the brain level and in seven functional systems defined by Yeo (Yeo et al. 2011). Considering the heterogeneity of the disorder, we repeated the same analyses on the subsamples of individuals with ASD divided into distinct symptoms profiles according to the total severity score and severity scores in the domain of social affect and restricted and repetitive behaviours domains.

### 4.1 Spatio-temporal changes in individuals with ASD compared to TD

At the global level, while no significant results were found between individuals with ASD and typically developed individuals, individuals with ASD more severely affected in the social affect domain globally showed a lower system diversity than the ones less affected in the SA domain. This pattern suggests a less mature functional repertoire of spatio-temporal connectivity patterns in participants with lower skills in social affect domain compared to individuals with less difficulties in this domain.

Changes in spatio-temporal dynamics turned out to be heterogenous across the different functional systems. When compared to the TD group, the ASD group showed significantly lower SD in the visual network while presenting a higher STD in the limbic functional system, the dorsal attention network and the visual network. Eye-tracking studies reported an increased idiosyncrasy on the neural level while watching dynamic social scenes related to lower scene understanding (Byrge et al. 2015) and higher presence of autistic symptoms (Bolton et al. 2020) which could relate to visual cortex alterations. The increased STD suggests more diversity and fluctuations in the functional patterns over time. This finding is consistent with the greater variability of functional pattens across time already observed in individuals with ASD (Falahpour et al. 2016).

Compared to other functional systems, the DM in the ASD group followed a specific behaviour with increased SD and decreased STD compared to the TD group. A specific SD and STD profile of DM network compared to other systems has already been observed in typical development (Vohryzek et al. 2019). The DM network has been associated with inner thinking, representation of self, and internal modes of cognition (Raichle et al. 2001) and plays an unique role in resting-state. Indeed, DM is a task negative system which anticorrelates in functional activity with other networks described as task-positive networks. Atypical DM function has already been reported in ASD (Padmanabhan et al. 2017). The dysfunction of this system is characterized by an overconnectivity within the nodes of the DM and underconnectivity with nodes outside of this system. The overconnectivity reported in static functional connectivity could be an indicator of the overrecruitment of heterogeneous functional system (higher system diversity) due to a boosted function of DM as a regulator of resting-state activity impacting cognitive flexibility.

At the same time, the decreased STD observed in DM suggest that the connectivity patterns involving the DM are more self-similar over time and could be related to longer dwell time in some states reported in dynamic functional connectivity analyses (Yao et al. 2016; Rashid et al. 2018). A few studies showed that the greater inter-individual variability across time in ASD was associated to a static global underconnectivity (Falahpour et al. 2016; Mash et al. 2019). In line with this idea, we can hypothesize that the overconnectivity observed in the DM could reflect a lower variability in the functional patterns over time. This corroborates the lower variability observed in the DM by He and al. (He et al. 2018).

### 4.2 Spatio-temporal connectivity changes along the symptoms dimension

Within the ASD group, SD and STD measures were able to distinguish between different profiles of symptoms. The group with most severely affected individuals in terms of total severity score and the group with predominant social affect domain symptoms showed similar SD and STD profiles. By definition, the total severity score (Gotham, Pickles, and Lord 2009) is based on social affect and RRB subscores. The social affect score has more items and a larger range possible values which could explain why the total severity score seems driven by the social affect symptoms. Lower system diversity was detected at the global level, and in the sensorimotor and the visual systems in individuals with ASD presenting more symptoms in social affect domain. This profile of symptoms (SA+) was also accompanied by a greater spatio-temporal diversity over all the functional systems and particularly the DM, the sensorimotor system and the ventral attention networks. These findings suggest that patients with more difficulties in social affect showed less integrative patterns (less spatially diverse patterns) and more fluctuations of the patterns over time. It supports the idea of a greater inter-individual variability across time (Falahpour et al. 2016; Mash et al. 2019) reflected by less socially engaged behaviour and sustained social attention.

Furthermore, the increased spatio-temporal diversity is stronger in the DM and the ventral attention network, strongly overlapping the salience network (Yeo et al. 2011), two of the cores networks involved in cognitive flexibility (Vinod Menon and Uddin 2010). Mash and al. (Mash et al. 2019) suggested that social communication deficits were more related to higher fluctuations mainly in language/salience network, in line with the higher spatio-temporal diversity we see in our ventral attention network. Overconnectivity between DM and central executive and underconnectivity between salience and central executive networks have already been reported in individuals with ASD (Abbott et al. 2016). The altered fluctuations observed in the current study could be a sign of failing its function of switching between central executive and DM networks. Other studies in dynamic functional connectivity support this hypothesis by highlighting less frequent transitions between salience-central executive/lateral frontoparietal and DM/medial frontoparietal in children with ASD (Marshall et al. 2020). These results encourage the idea of disruptions in brain mechanisms involved in cognitive flexibility in individuals with ASD. Future work is crucially needed to further investigate the spatio-temporal dynamic of cognitive flexibility in individuals with ASD.

Individuals with severe RRB symptoms showed a different spatio-temporal dynamic profile compared to the patients whose profile was characterized by predominantly severe symptoms in social affect. These patients showed a lower spatio-temporal diversity in the DM and a trend to lower STD in other functional systems as well but the FP. From a dynamic point of view, Mash and al. (Mash et al. 2019) already showed that the RRB subscales may be higher for individuals who spend longer time in each state and switch less frequently between states. This is consistent with the idea of less diversity in the fluctuations over time, especially in the DM network, that we observed in the current study for the most severe RRB group. Finally higher levels of RRB have been associated with cognitive inflexibility (Dajani and Uddin 2015). Further analyses should confirm that lower STD in the DM for patients with severe RRB scores could be the expression of the altered mechanisms of cognitive flexibility. Finally, it has been shown that subtypes of RRB had different profiles in connectivity with stereotyped behaviours (i.e. purposeless movements repeated in a similar manner) more associated to positive functional connectivity in dorsal attention and subcortical regions while restricted behaviours (i.e. limited interest, activity or focus) (Lam and Aman 2007) were more related to the connectivity between DM and dorsal attention for example (McKinnon et al. 2019). A dynamic perspective on these different subtypes could help to highlight more alterations between these subtypes.

The opposing pattern of the spatio temporal dynamic between different symptoms profile highlights the necessity of consideration of tremendous heterogeneity of the disorder. Our results suggest that the two main symptom dimensions in ASD (SA and RRB) may be depending on the complex and somewhat opposing patterns of the underlying brain dynamics. While the social communication deficits might be driven by the highly temporally variable patterns of brain activity involving DMN and SAL network, a more conservative, less variable DMN pattern was seen in patients with more symptoms in RRB domain.

### 4.3 Methodological considerations and limitations

System and spatio-temporal diversity are based on the connected components extracted from the multilayer graph generated with the spatio-temporal connectome (Griffa et al. 2017). This framework combines functional co-activations from framewise analysis on BOLD signal time series and an average structural connectome. The usage of an average connectome reduces the bias of the existence of false positive prominent in an individual structural connectivity matrix (Zalesky et al. 2016). However, taking an average structural connectome also means the information we integrate in this framework is not specific to the participants’ dataset. Including the information specific to an individual would allow to really merge structural and functional information at the subject level. For now, with the usage of an average connectome, we investigated functional dynamics supported by direct structural pathways. These measures are group-level measures and are not defined at the subject level. This group definition limits the possibility of correlations with demographic information or behavioural or clinical scores.

## 5. CONCLUSIONS

We characterized the spatio-temporal brain dynamics in individuals with ASD (using ABIDE 1 database) using two novel measures, namely the spatio-temporal diversity and the system diversity. On the one hand, individuals with ASD showed lower integration of various cognitive systems within brain processes (lower system diversity) and a less stable functional activity with less self-similar brain processes over time (higher spatio-temporal diversity). On the other hand, DM network exhibited an opposite profile of spatio-temporal dynamics, consistent with previous work reporting a distinctive pattens in static connectivity as well. The decrease in system diversity and the increase in spatio-temporal diversity were stronger in patients presenting more severe symptoms, particularly in case of prominent social affect symptoms, supporting the idea that these alterations could be related to the cognitive deficits in ASD. The increased patterns of stability within the DM (lower spatio-temporal diversity) were characteristic of individuals with prominent RRB symptoms suggesting the existence of inflexible cognitive processes specific to these behaviours. These results showed first evidence that distinct profiles of spatio-temporal dynamic exist along the symptoms dimension.

## ABBREVIATIONS

ASD: Autism Spectrum Disorder
CTRL: controls
SD: system diversity
STD: spatio-temporal diversity
ABIDE: Autism Brain Imaging Data Exchange
CCs: Connected Components
FD: framewise displacement
DMN: default mode network
VIS: visual
FP: frontoparietal
DA: dorsal attention
VA: ventral attention
SM: somatomotor
LIMB: limbic
RRB: restricted and repetitive behaviours

## ACKNOWLEDGEMENTS

This study was supported by the National Center of Competence in Research “Synapsy,” financed by the Swiss National Science Foundation (SNF, Grant Number: 51NF40_185897), by SNF Grants to MS (#163859 and #190084). Funding sources for the datasets comprising the 1000 Functional Connectome Project are listed at fcon_1000.projects.nitrc.org/fcpClassic/FcpTable.html. Funding sources for the ABIDE dataset are listed at fcon_1000.projects.nitrcc.org/indi/abide.

